# Transformations, Lineage Comparisons, and Analysis of Down to Up Protomer States of Variants of the SARS-CoV-2 Prefusion Spike Protein Including the UK Variant B.1.1.7

**DOI:** 10.1101/2021.02.09.430519

**Authors:** Michael H. Peters, Oscar Bastidas, Daniel S. Kokron, Christopher E. Henze

## Abstract

Monitoring and strategic response to variants in SARS-CoV-2 represents a considerable challenge in the current pandemic, as well as potentially future viral outbreaks of similar magnitude. In particular mutations and deletions involving the virion’s prefusion Spike protein have significant potential impact on vaccines and therapeutics that utilize this key structural viral protein in their mitigation strategies. In this study, we have demonstrated how dominant energetic landscape mappings (“glue points”) coupled with sequence alignment information can potentially identify or flag key residue mutations and deletions associated with variants. Surprisingly, we also found excellent homology of stabilizing residue glue points across the lineage of *β* coronavirus Spike proteins, and we have termed this as “sequence homologous glue points”. In general, these flagged residue mutations and/or deletions are then computationally studied in detail using all-atom biocomputational molecular dynamics over approximately one microsecond in order to ascertain structural and energetic changes in the Spike protein associated variants. Specifically, we examined both a theoretically-based triple mutant and the so-called UK or B.1.1.7 variant. For the theoretical triple mutant, we demonstrated through Alanine mutations, which help “unglue” key residue-residue interactions, that these three key stabilizing residues could cause the transition of Down to Up protomer states, where the Up protomer state allows binding of the prefusion Spike protein to hACE2 host cell receptors, whereas the Down state is believed inaccessible. Thus, we are able to demonstrate the importance of glue point residue identification in the overall stability of the prefusion Spike protein. For the B.1.1.7 variant, we demonstrated the critical importance of D614G and N5017 on the structure and binding, respectively, of the Spike protein. Notably, we had previously identified D614 as a key glue point in the inter-protomer stabilization of the Spike protein prior to the emergence of its mutation. The mutant D614G is a structure breaking Glycine mutation demonstrating a relatively more distal Down state RBD and a more stable conformation in general. In addition, we demonstrate that the mutation N501Y may significantly increase the Spike protein binding to hACE2 cell receptors through its interaction with Y41 of hACE2 forming a potentially strong hydrophobic residue binding pair. We note that these two key mutations, D614G and N501Y, are also found in the so-called South African (SA; B.1.351) variant of SARS-CoV-2. Future studies along these lines are, therefore, aimed at mapping glue points to residue mutations and deletions of associated prefusion Spike protein variants in order to help identify and analyze possible “variants of interest” and optimize efforts aimed at the mitigation of this current and future virions.

## I. Introduction

*β* coronaviruses represent one (B) of four genera (A,B,C,and D) of RNA positive sense viruses in the Nidovirales order [1, 2]. The current pandemic COVID-19 caused by SARS-CoV-2 is the latest in human viral outbreaks of this genera being preceded by the Middle Eastern Respiratory Coronavirus (MERS-CoV) and the SARS-CoV outbreak of 2002. SARS-CoV-2 continues to exhibit high rates of transmission and infection across the globe in the current pandemic. Of great present concern are variants that may show relative increased transmission and infection rates and, in addition, may present challenges to current and developing vaccines as well as therapeutics aimed at mitigation of this deadly virion. Of note is that this virus has a genome size of ∼ 30 kilobases and an intrinsic proofreading mechanism to reduce mutation rates [3]. The mutation rate of SARS-CoV-2 has been estimated to be ∼ 10^−3^ substitutions per site per year [3]. Genomic sequences of SARS-CoV-2 continue to be deposited in the GSAID (Global Initiative on Sharing all Influenza Data), which has allowed for the study of structural implications of mutations [4]. For example, the Spike protein mutation D614G has been associated with higher upper respiratory tract viral loads and appears to be omnipresent in recent genomic sequences across the globe [4, 5, 6]. In addition, another variant called the UK Variant or VOC 202012/01 or B.1.1.7 (classification system [7]) has been identified as a highly transmittable variant and involves both deletions and mutations in the Spike protein of this virion, including D614G. So, it is of great importance to determine how current and future variants may translate into altered transmission rates, viral loading differences, antibody and vaccine escape, and resistance to currently developing therapeutics. Here we focus on the analysis of mutations of the prefusion Spike protein, due to its importance to vaccines and therapeutics, as a partial guide to the potential effects of its mutations on structure, function, and possible behavioral changes of this virion.

A distinct characteristic of the coronaviruses are their large, trimeric Spike proteins that densely decorate the virion surface [8, 9, 10]. The Spike protein consists of three homologous protomers or chains where each one is ∼ 1200 amino acid residues in length (Fig. 1). In its prefusion state, each protomer consists of two large domains called S1 (most distal from the virion membrane) and S2 (most proximal to it’s membrane). In general, the S1 domain represents a prefusion domain (∼ 650 residues) and the S2 domain (∼ 600 residues) is the fusion domain. In general, the S2 or fusion machinery domain is relatively rigid with strong noncovalent intra and interchain interactions facilitated by helical secondary structures, whereas the S1 domain, which contains the host cell receptor binding domain (RBD) and N-terminal domain (NTD) in a V-shaped configuration (Fig. 1), is weaker, flexible, and characterized by beta-strand secondary structural motifs [11]. We note that the S1 domain of the Spike protein is shed in the transition from the prefusion state to the fusion state. The configuration of the RBD in the prefusion state is further characterized as being in the so-called “Up-state” or “Down-state” depending on the position of the RBD relative to the center of mass of the prefusion Spike protein. For example, in the Up-state, the RBD of both SARS-CoV and SARS-CoV-2 is more exposed and able to bind to its ACE2 (Angiotensin Converting Enzyme 2) receptor on the surface of human epithelial cells (Type I and II pneumocytes; also, alveolar macrophage and nasal mucosal cells), but in the “Down-state” the RBD is believed to be more hidden and significantly reduced to ACE2 binding and to cellular infection [12, 13, 14]. Quantified structural comparisons of the RBD configuration in the Up versus Down protomer states of SARS-CoV-2 have recently been done that include angular positions of the RBD relative to the NTD of a given protomer [8]. Henderson et al [13] also quantified angular differences in the S1 subdomains of the Spike protein across the *β* coronaviruses SARS-CoV-2, SARS-CoV, MERS and HKU1 and developed mutational forms that can alter the equilibrium of Up versus Down states.

**Figure 1:**
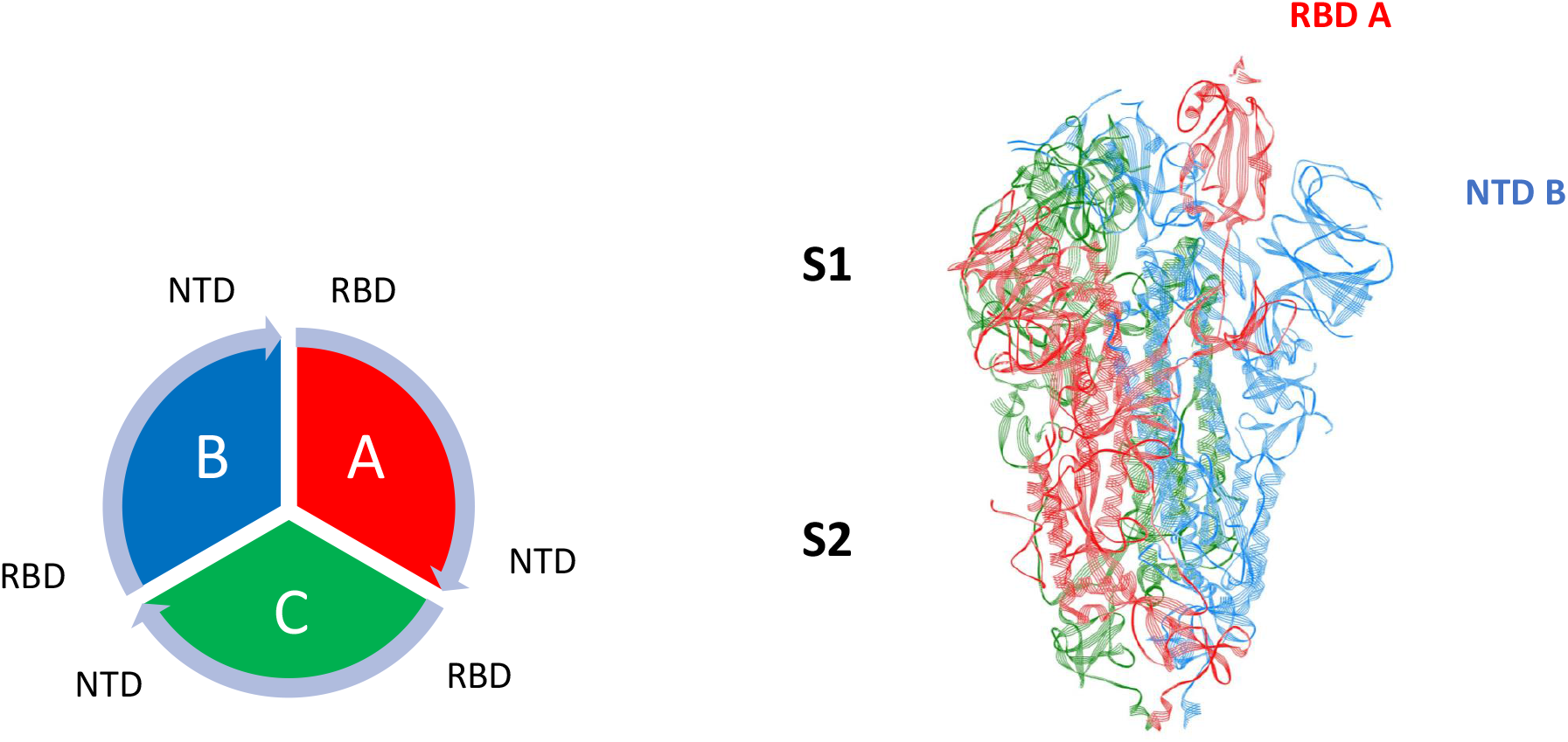
SARS-Cov-2 *β* coronavirus (PDB ID: 6VSB) with one Up (A) and two Down (B,C) showing the S1 (binding ectodomain) and S2 (fusion) domains. Also shown is the overall chain interaction configuration looking at the trimer from the top view.

Given the critical importance of emerging variants of SARS-CoV-2 to vaccines and therapeutics, it is important to analyze the effects of mutations and/or deletions on the stability and dynamics of the Spike protein. Previously, we studied the stability and dynamics of the entire Spike protein of SARS-CoV-2 using a combination of all-atom dominant energetic analyses and biophysical computational molecular dynamics using published structures of the trimeric Spike protein [11]. We determined energetically dominant, non-covalent intra-protomer and interprotomer interactions, called “glue” points or “hot” spots that help stabilize the entire trimeric protein structure. For example, we previously identified D614 as a key glue point with neighboring protomer residues ([11, 15], Table S1) prior to its emergence in current variants. We also mutated a key hot spot (‘latch’ residues) associated with intra-protomer interactions in order to demonstrate the ability for single protomers to change from Down to Up states. However, it was further demonstrated that in complete trimeric structures such transitions are held in check by inter-protomer interactions, specifically, the RBD of any protomer with the NTD of its neighbor.

In the current study, we computationally analyze structure and dynamics of key mutations associated with Down versus Up protomer states of SARS-CoV-2 prefusion Spike protein; in particular, we examine a theoretically-based triple mutant, based on our glue point mappings, that is shown to destabilize neighboring RDB-NTD interactions and lead to a transition of Down to Up protomer states within the complete trimeric protein. We are also interested in how these stabilzing RBD-NTD interactions may be conserved across the lineages of *β*-corona viruses SARS-CoV, MERS-CoV and HCoV, since such comparisons may be useful in identifying critical deletions/mutations across current and future variants of SARS-CoV-2 and possibly future emerging *β*-coronaviruses. We demonstrate through sequence alignment and energetic mappings highly conserved stabilizing glue point residues across the lineages of *β*-coronaviruses. In addition, using these tools, we critically examine the UK variant B.1.1.7, including D614G mutation, in order to discern key differences in protomer configurations that could potentially impact vaccine and therapeutic efforts aimed at the debilitation of the current pandemic. We then dynamically analyze any associated conformational changes using all-atom molecular dynamics of the trimeric prefusion spike protein of B.1.1.7 over a 0.5 microsecond time period in order to ascertain key differences in behavior from the wild type. We are able to demonstrate dynamic changes in the B.1.1.7 spike protein that can be traced to two key mutations resulting in a more accessible RBD (D614G) and simultaneously stronger binding to ACE2 (N501Y). These key mutations are also present in the rapidly emerging South African variant B.1.351. Our findings and analysis may have general applicability and may be important at ascertaining the potential effects of future variations of this virion on vaccine and therapeutics, as well as possible relations to infection and transmission rates, in an attempt to stay ahead via structure→function analysis of emerging variants. We note that our analysis here is focused on the prefusion state of the Spike protein and does not consider the additional steps of fusion, uptake, and virion replication in host cells, which are all important to the overall transmission and infection process.

## Materials and Methods

### Molecular Dynamics

Explicit solvent molecular dynamics (MD) simulations of the novel coronavirus Spike protein were performed using the NAMD2 program [16]. We used the CHARMM-Gui [17] with the CHARMM36m force field along with TIP3P water molecules to explicitly solvate proteins and add any missing residues from the experimental structure files. Disulfide bonds and glycosylated sites were all included. Simulations were carried out maintaining the number of simulated particles, pressure and temperature (the NPT ensemble) constant with the Langevin piston method specifically used to maintain a constant pressure of 1 atm. We employed periodic boundary conditions and initial equilibration for a water box simulation volume as well as the particle mesh Ewald (PME) method with a 20 Å cutoff distance between the simulated protein and water box edge. The integration time step was 2 femtoseconds with our protein simulations conducted under physiological conditions (37 C, pH of 7.4, physiological ionic strength with NaCl ions, LYS and ARG were protonated and HIS was not). All mutations were added via the CHARMM-Gui [17] and for deletions we chose to use Glycine as a structure breaking mutation in lieu of deleting residues, since exact structural information on deletions is currently lacking. Any other methods to revise structure, such as the SWISS model [18], would still be approximate and not based on the actual protein folding dynamics, whereas the more straight-forward, structure breaking Glycine represents a good test of the potential role of *deletions* on structure, as will be demonstrated here. All MD results given here were also repeated several times in order to help confirm trends in data.

### Sequence Alignment

Multiple sequence alignment was preformed using Clustal Omega [19]. Clustal Omega uses a structure guided hidden Markov model (HMM) for multiple sequence alignment. Sequences were obtained directly from the PDB files across four different *β* corona viruses: SARS-CoV (6ACD) [14], SARS-CoV-2 (6VSB) [10], MERS-CoV (6Q04) [20], and HCoV (6OHW) [21]. Output format was selected as ClustaW with character counts.

### All-Atom Energetic Mappings

Previously [11, 15], we analyzed the complete inter-protomer and intra-protomer interactions across two independently published structure files (6VSB and 6VYB) for SARS-CoV-2 trimeric Spike protein using the open source energy mapping algorithm developed by Krall et al [22]. This spatial and energetic mapping algorithm efficiently parses the strongest or most dominant non-covalent atom-atom interactions (charge and partial atomic charge, Born, and van der Waals forces), according to empirically established parsing criteria, based on the *ab initio* AMBER03 force field model. Following our previous studies, the parsing criteria were taken as the upper limit of −0.1*kT* units for Lennard-Jones (van der Waals) criteria and −0.3*kT* units for Coulombic interactions, although lower values can also be specified in the analysis part of the mappings in order to further refine the results [22]. Note that in the all-atom analysis dominant van der Waals interaction forces are commonly associated with nonpolar atom-atom interactions and hydrophobic protein interaction regions, whereas the Coulombic partial charge and charge interactions are commonly associated with hydrophilic protein interaction regions and can include hydrogen bonding and backbone atom partial charge interactions.

### RMSF and Hinge Angle Determinations

Here we follow the recent hinge angle designation of Peng et al [23] in order to help quantify and compare Up versus Down protomer states, namely ∠ ASP406-VAL991-ALA622. Note that the vertex selected (VAL991) is in the rigid S2 domain and therefore is approximately fixed in the body frame of the protein. This designation helps to correct for any so-called “tumbling” effects associated with translation and rotations of the center of mass of the protein over large time scales necessary for these types of simulations. Those authors further designated hinge angles in the range 52.2 deg to 84.8 deg as ACE2 accessible domains (Up states) and angles in the range 31.6 deg to 52.2 deg as ACE2 inaccessible (Down states). Here, we also define the “Arm” and “Leg” of the hinge angle as the C-*α* distance between *ASP* 406 − *V AL*991 and *V AL*991 − *ALA*622, respectively.

Root Mean Square Fluctuations (RMSF) C-*α* values across the 1124 residues for any protomer were determined according to

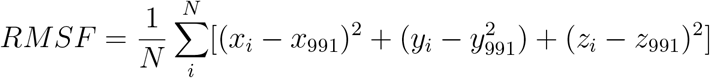

where (x,y,z) are the cartesian coordinates of any C-*α* residue, N is the number of snapshots considered, and deviations are measured relative to the body frame or VAL991 for consistency with hinge angle calculations and to correct for tumbling effects. Here we take snapshots of structures after every 1.0 nsec.

## Results

### Sequence Alignment and Glue Point Residues Across Lineages of *β* coronaviruses

The color map sequence alignment across the entire Spike protein for the four *β* coronaviruses as obtained from Clustal Omega is shown in Fig. 2. As can be seen the greatest overall alignment homology is with the S2 or fusion domain and S1-NTD of this protein, and the greatest variation is in the RBD of S1. We also mapped from the original structure files the dominant energetic contacts or glue points of the stabilizing RBD-NTD neighboring chain interactions across these lineages as shown in Fig. 4, where we have superimposed predicted glue points over the sequence alignment. Somewhat surprisingly we found excellent sequence homology across almost all glue points despite clade differences among these lineages; below we refer to these as “sequence homologous glue points”. Additionally, within the SARS-CoV and SARS-CoV-2 clade we found larger numbers of atom-atom interactions for the same glue point residues associated with SARS-CoV, indicative of a much stronger and more stable Down state configuration (Fig. 5). Interestingly, SARS-CoV-2 demsonstrates the least total number of glue points and associated stability across these lineages. Note that these results are based on independently obtained experimental PDB deposited structure files: SARS-CoV-2: 6VSB [9], 6VYB [10]; SARS-CoV: 6ACD [14], 6CRZ [24], as shown in Fig. 5.

**Figure 2:**
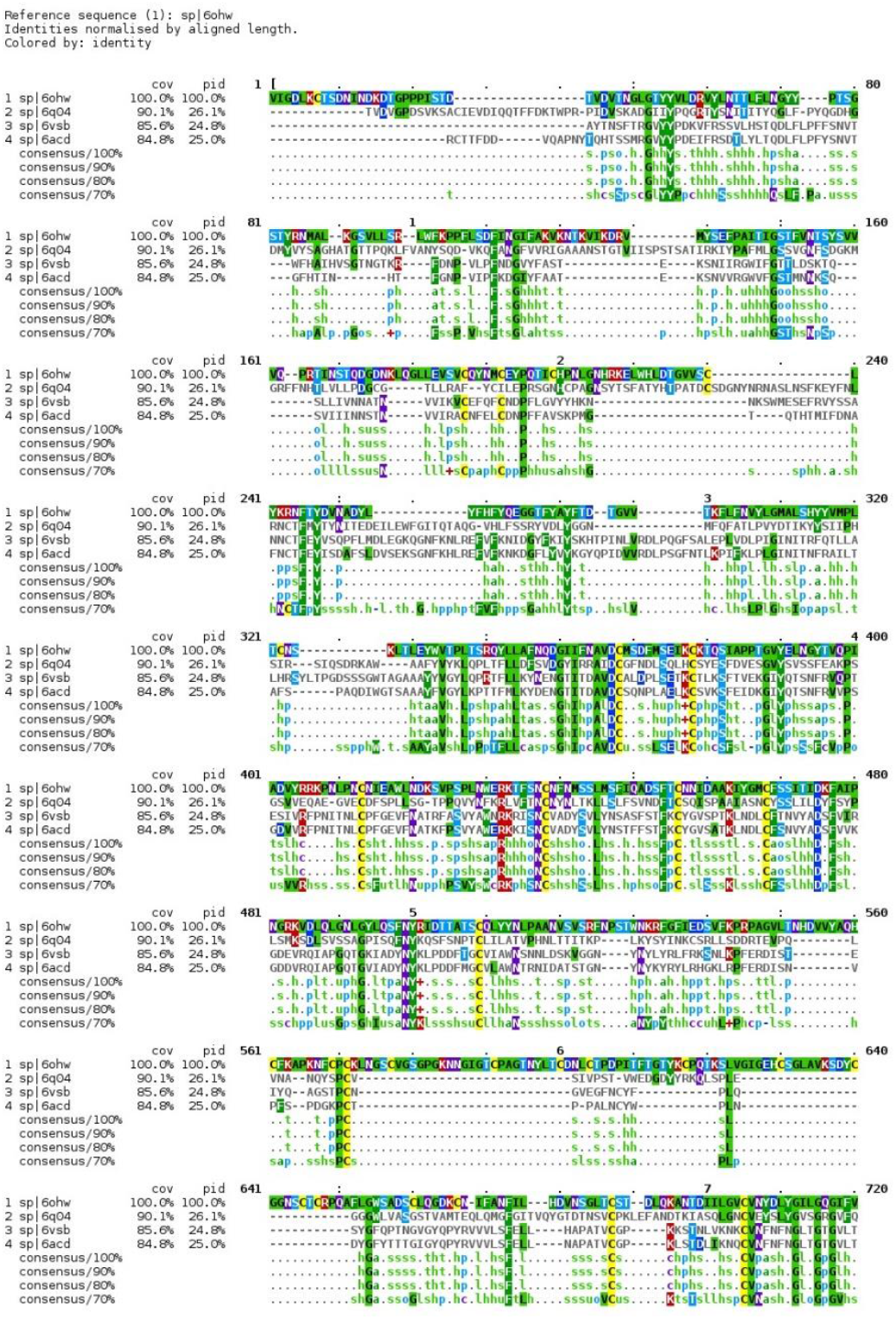
Sequence alignment for SARS-Cov, SARS-CoV-2, MERS, and HCoV. Original .ppt file is included in the Supplementary material for ease of viewing.

**Figure 4:**
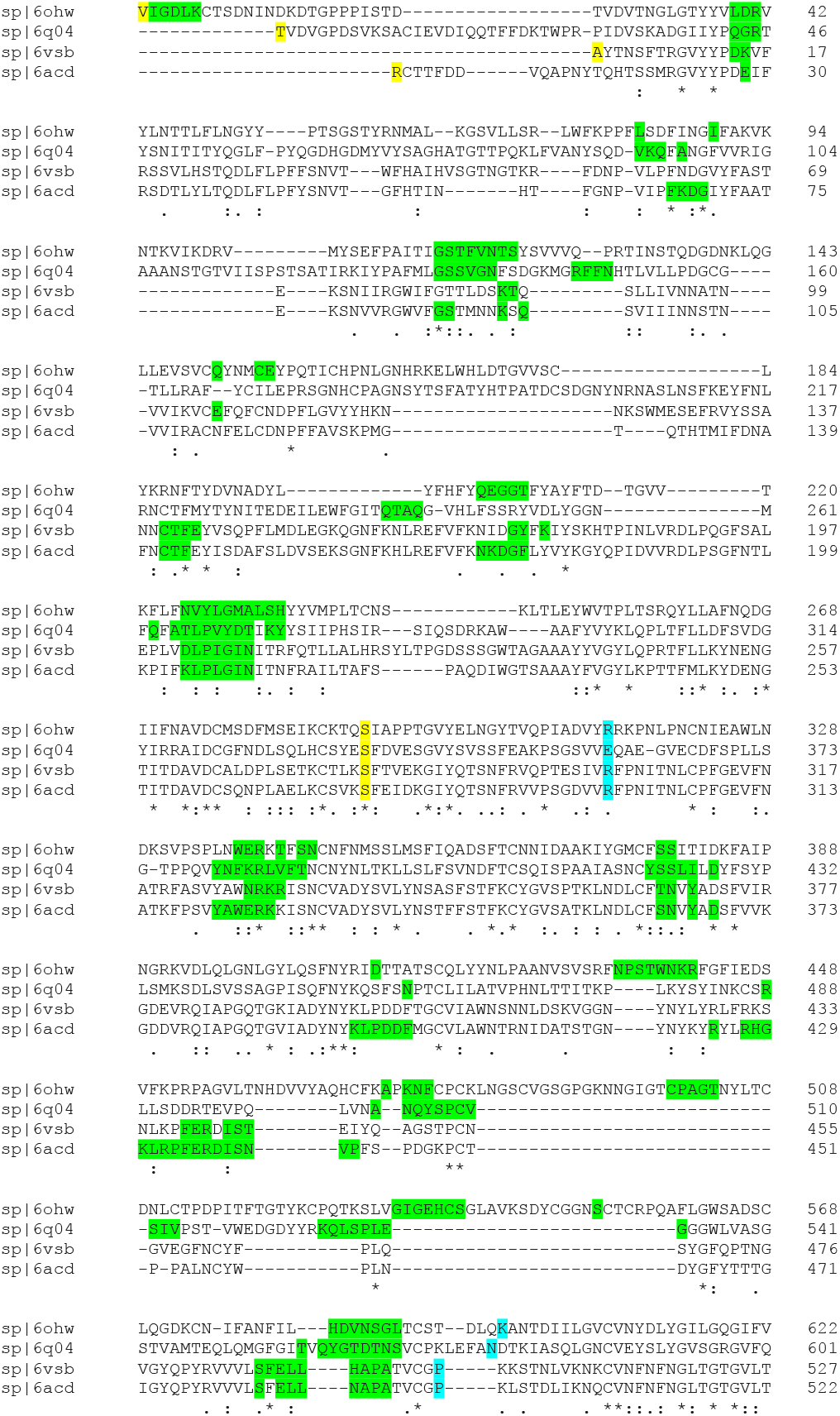
Combination of the Sequence Alignment Map with the Glue Point Map for the S1 Domain Across the *β* coronaviruses. Green shaded letters are the residues associated with dominant energetic enteractions or glue points between the RBD and its neighboring NTD in the Down state. Yellow shaded letters mark the start and end of the NTD and blue shaded letters mark the beginning and end of the RBD across these lineages.

**Figure 5:**
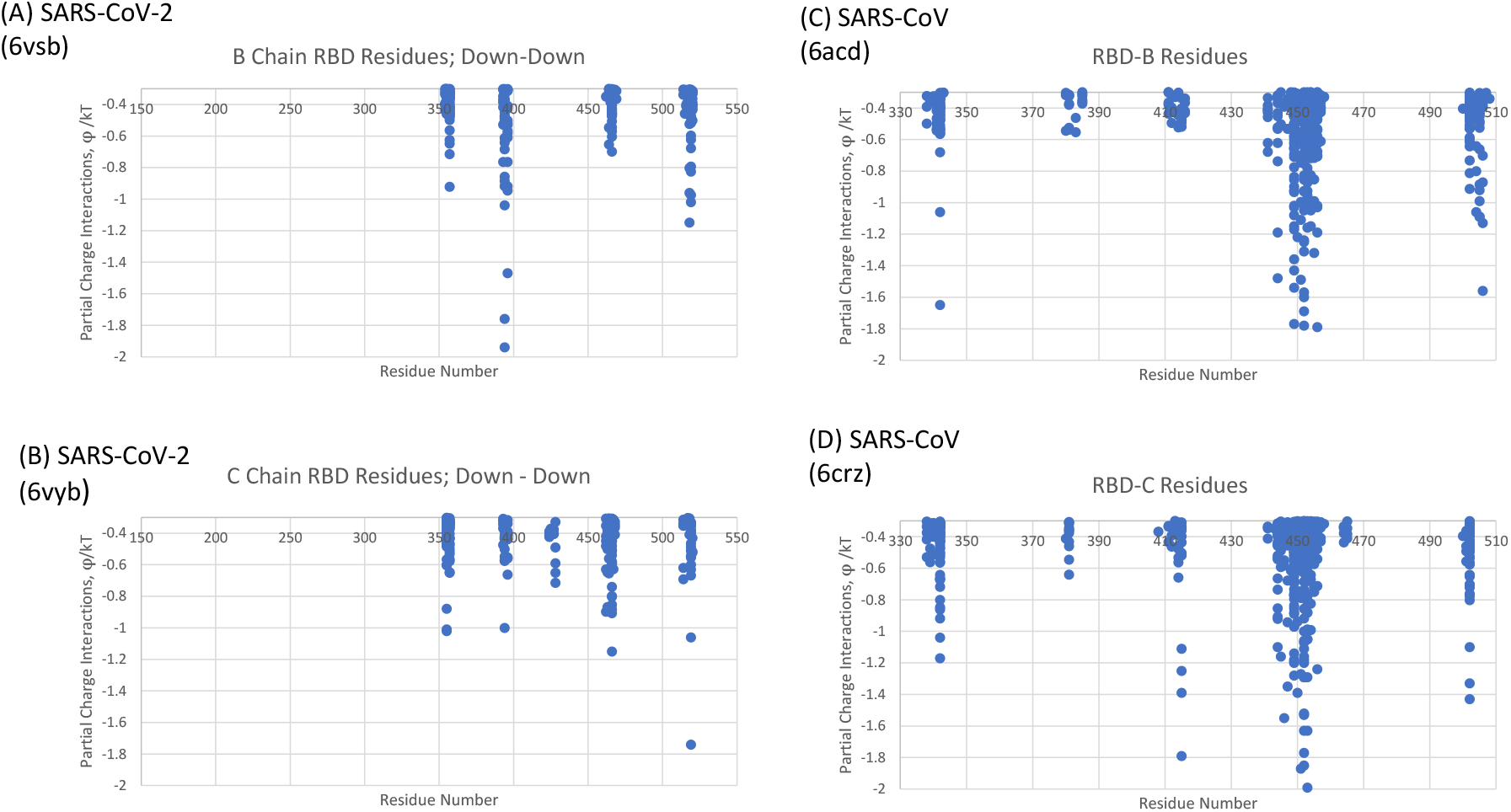
Comparison of SARS-CoV, (C) and (D), and SARS-CoV-2, (A) and (B), RBD-NTD neighboring chain glue points. Note that all protomers are in the Down state here. See [11] Supplementary Data for (A) and (B); Data for (C) - Table S1; Data for (D) - Table S2

### Triple Mutant versus Wild Type

Previously, and as partially shown in Fig. 5 for Down-Down state interactions, we identified three critical glue point residues that help stabilize RBD-NTD inter-protomer interactions across both Down-Down and Up-Down states of SARS-CoV-2: viz., ARG357, ASN394, and HIS519 ([11], Figs. 3 and 4). These interactions helped prevent “latch” release from Down to Up states associated with Down state intra-protomer latch residues: GLN564 to ALA520-PRO521-ALA522. Note from Fig. 4 that these three stabilizing residues are also part of the sequence homologous glue points across the lineages of *β*-coronaviruses. Here we examine the triple Alanine mutant ARG357ALA, ASN394ALA, and HIS519ALA in order to determine if these key glue points alone could cause a conformational change in the absence of any latch mutations. Note that this so-called “Alanine screening” should diminish side chain interactions of those residues (“unglue”) without significant initial structure changes. Figure 6 A-D shows MD calculated values of the hinge angle and RMSF values for what we have called wild type (WT) SARS-CoV-2 and the theoretical triple mutant. It is clear that a longer time period of 150 nsec is required to reach a dynamic equilibrium state of either of these proteins from the starting configurations that include an initial NVT equilibrium period. For fairness, we determined the RMSF values after the first 150 nsec of simulation. The WT hinge angle equilibrates at 1 radian or approximately 60 deg putting it on the lower end of the ACE2-accessible region according to the criteria of Peng et al. [23] Additionally, the theoretical triple mutant Down B Chain transitions to the Up Chain state after approximately 220 nsec (Fig. 6C) and continues the Up State transition as a result of these key residue mutations (Supplementary Data S1). The triple mutant also shows more flexibility in its S1 domain, according the calculated RMSF values, as compared to WT as expected for the negation of its three key stabilizing, glue point residues. Note that the RMSF values for the S2 domain are the same between WT and the theoretical triple mutant as expected. These results show how sequence information superimposed on glue point maps followed by biophysical computations can help guide and understand the possible outcomes of mutations to protein function.

**Figure 3:**
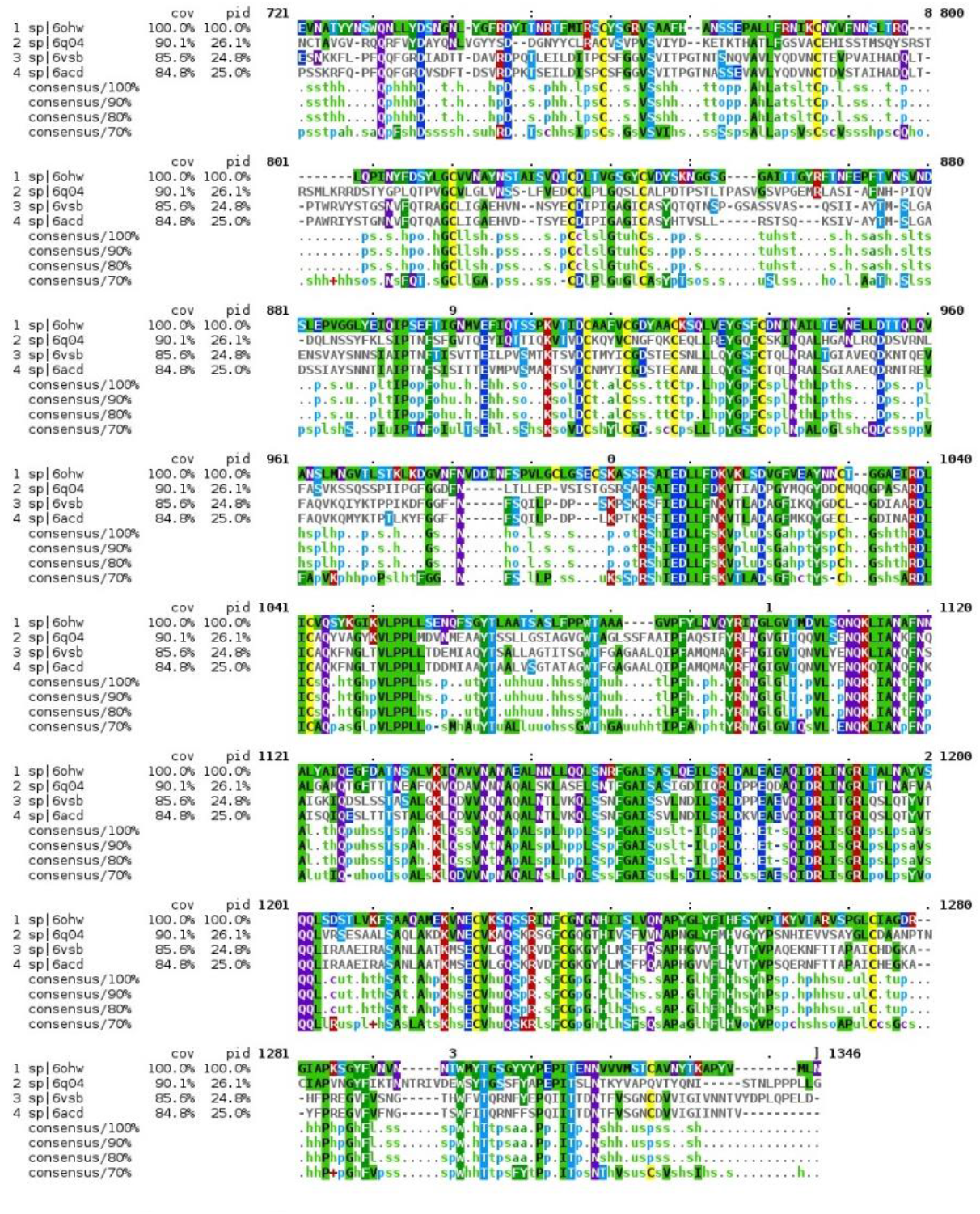
Sequence alignment for SARS-Cov, SARS-CoV-2, MERS, and HCoV. Original .ppt file is included in the Supplementary material for ease of viewing.

**Figure 6:**
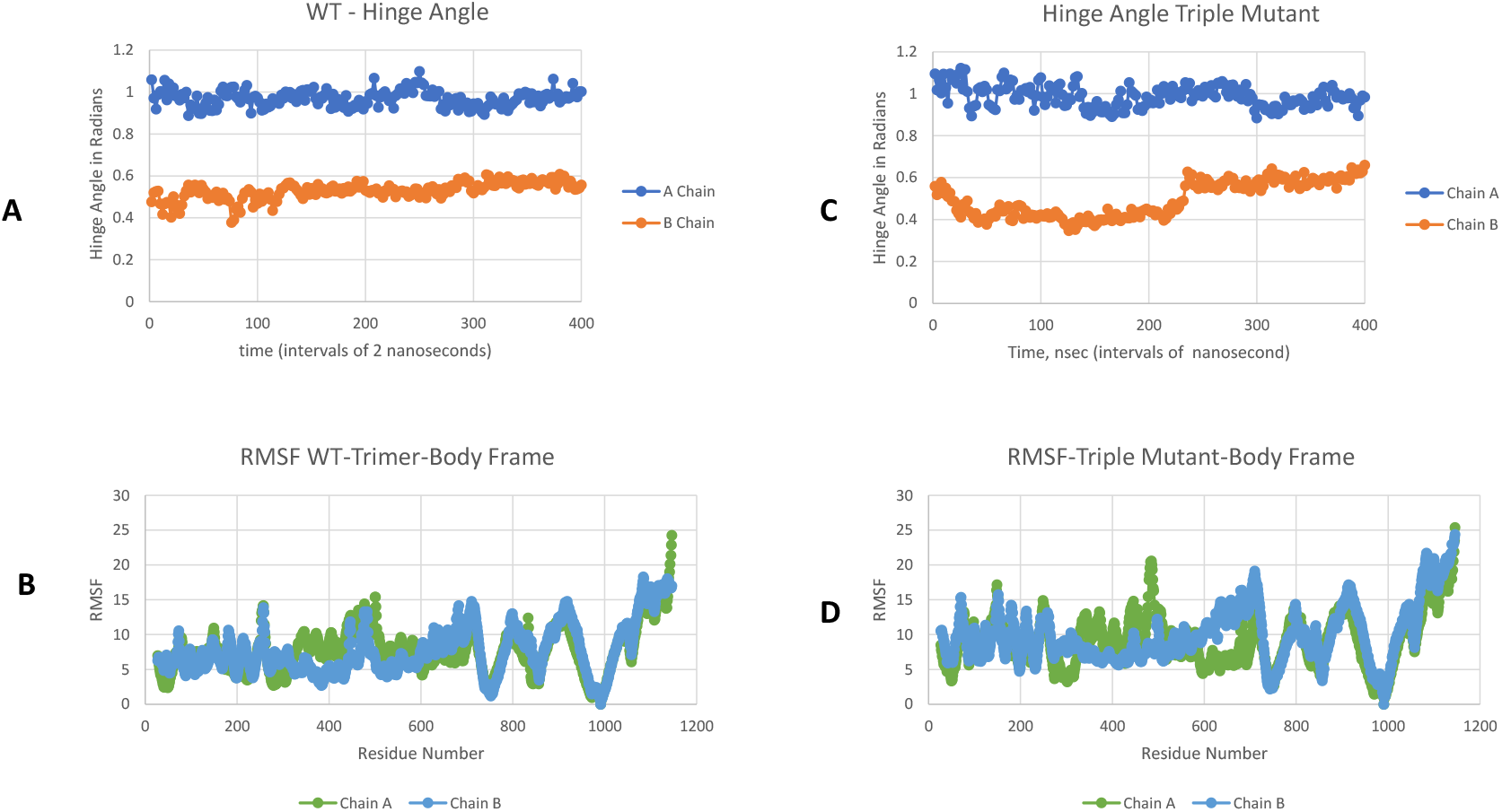
Hinge angle and RMSF values for Wild Type (A) and (B) frames and Triple Mutant and (D) frames, respectively; Chain A is Up and Chain B is Down. RMSF values shown for the last 200 nsec of the simulation.

### UK Variant B.1.1.7

A summary of the mutations and deletions of the UK Variant B.1.1.7 are given in Table 1. Also shown are the partner glue point residues predicted by OpenContact when the B.1.1.7 residues are in the Up state and mapped to a neighboring Down state protomer or in the Down state and mapped to a neighboring Down state protomer. As can be seen only A570D and D614G involve glue point partners within the Spike protein. None of the glue point partners are involved in the NTD-RBD sequence homologous regions presented previously. Additionally, we mapped hACE2 binding of SARS-CoV-2 RBD according to the full-length hACE2 structure file PDB ID: 6M17 [12]. We have overlayed the dominant glue point residues to hACE2 in red as shown in Fig. 7. The residue N501 is a key binding partner to hACE2 (Supplementary Table S3.) and this includes conspicuously strong interactions with Y41 of ACE2. The N501Y mutation may potentially increase binding to hACE2 significantly due to the highly favorable Y-Y hydrophobic interaction pair in the new mutant state (Table S3), however this remains to be concretely verified. Note that both D614G and N501Y are also present in the South African variant (B.1.351).

**Table 1:** Summary of Mutations and Deletions of B.1.1.7 Variant

| Mutation/Deletion | Up → Down | Down → Down | NTD-RBD Homologous | ACE2 Binding |
| --- | --- | --- | --- | --- |
| N501Y | - | - | No | Yes |
| A570D | S2: 960-967 | S2: 963-967, 875, 1000 | No | No |
| D614G | S2: 854-860 | S2: 733-735, 854-861 | No | No |
| P681H | - | - | No | No |
| Delete V69-S70 | - | - | No | No |
| Delete Y144-Y145 | - | - | No | No |

**Figure 7:**
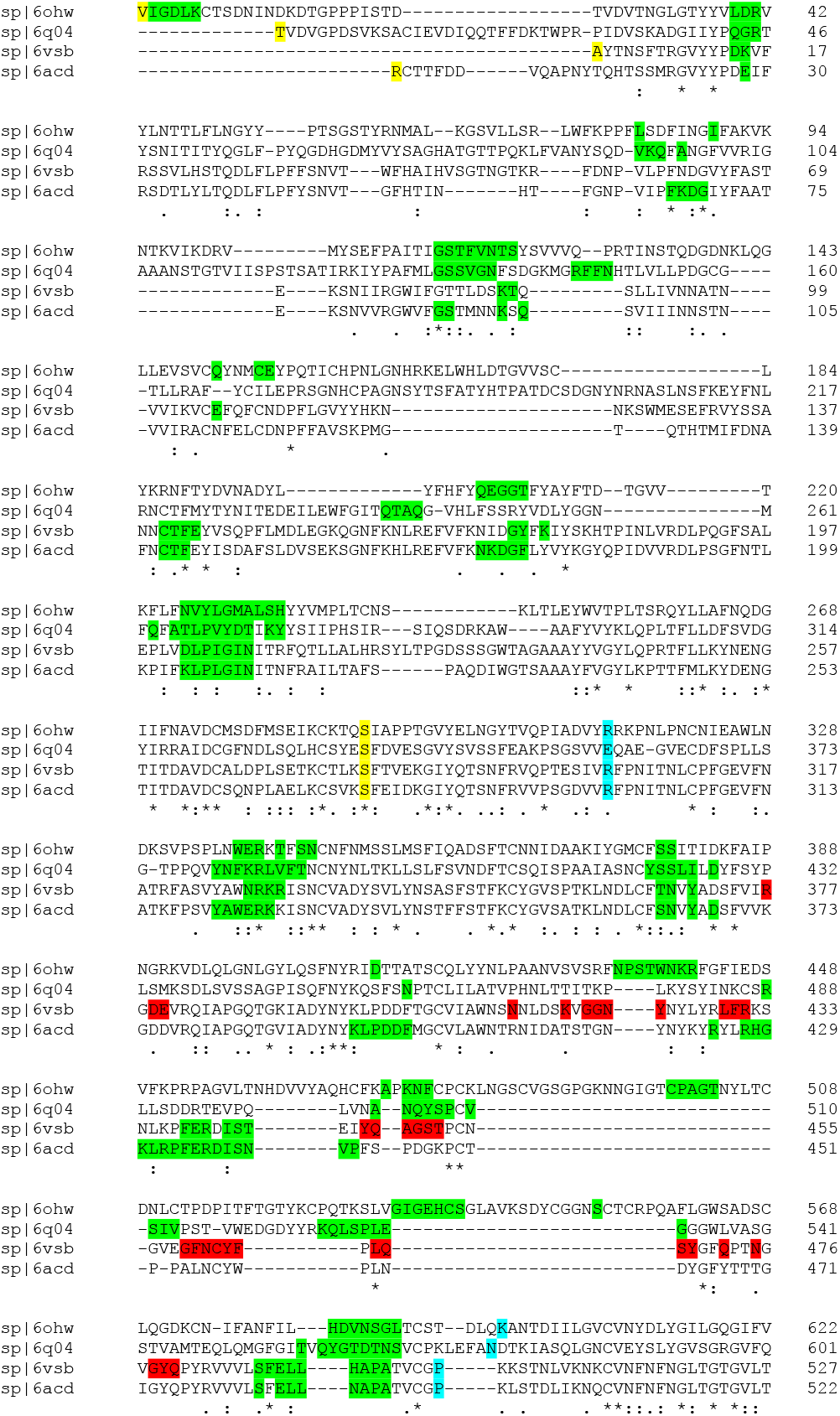
SARS-CoV-2 Binding residues to hACE2 (red shaded letters) shown on the combination of the Sequence Alignment Map with the Glue Point Map for the S1 Domain Across the *β* coronaviruses. See Table S3 for completer data.

Next, we performed long time MD simulations of B.1.1.7 as described in the Methods section and shown in Fig. 8 (cf. Fig. 6). As can be seen B.1.1.7 variant stabilizes to the WT Up (Chain A) and Down (Chain B) hinge angles after approximately 150 nsec of simulation. Because of the lack of significant change of hinge angles observed, we looked more closely at the specific length measures associated with this angle, i.e., the Arm and Leg lengths discussed in the Methods section. Figure 9 shows the results of Arm and Leg calculations across each of the Spike proteins: WT, Triple Mutant and UK Variant. The Leg lengths associated with the more proximal S1 domain to S2 showed very little differences across all three variants; although, again, B.1.1.7 demonstrated more stability. Similarly, the Up chain Arm lengths also showed little changes between variants with B.1.1.7 again demonstrating better stability. Interestingly, however, the Down chain Arm lengths showed conspicuous differences, where both the Triple Mutant and UK Variant demonstrated a longer “reach” as compared to WT. Note that deletions associated with B.1.1.7 (Table 1.)and modeled by structure breaking Glycine mutations here, showed increased flexibility of the NTD, as expected, but did not show any conspicuous differences outside of that domain. (Also, see Supplementary Movies: UKMutvsWT)

**Figure 8:**
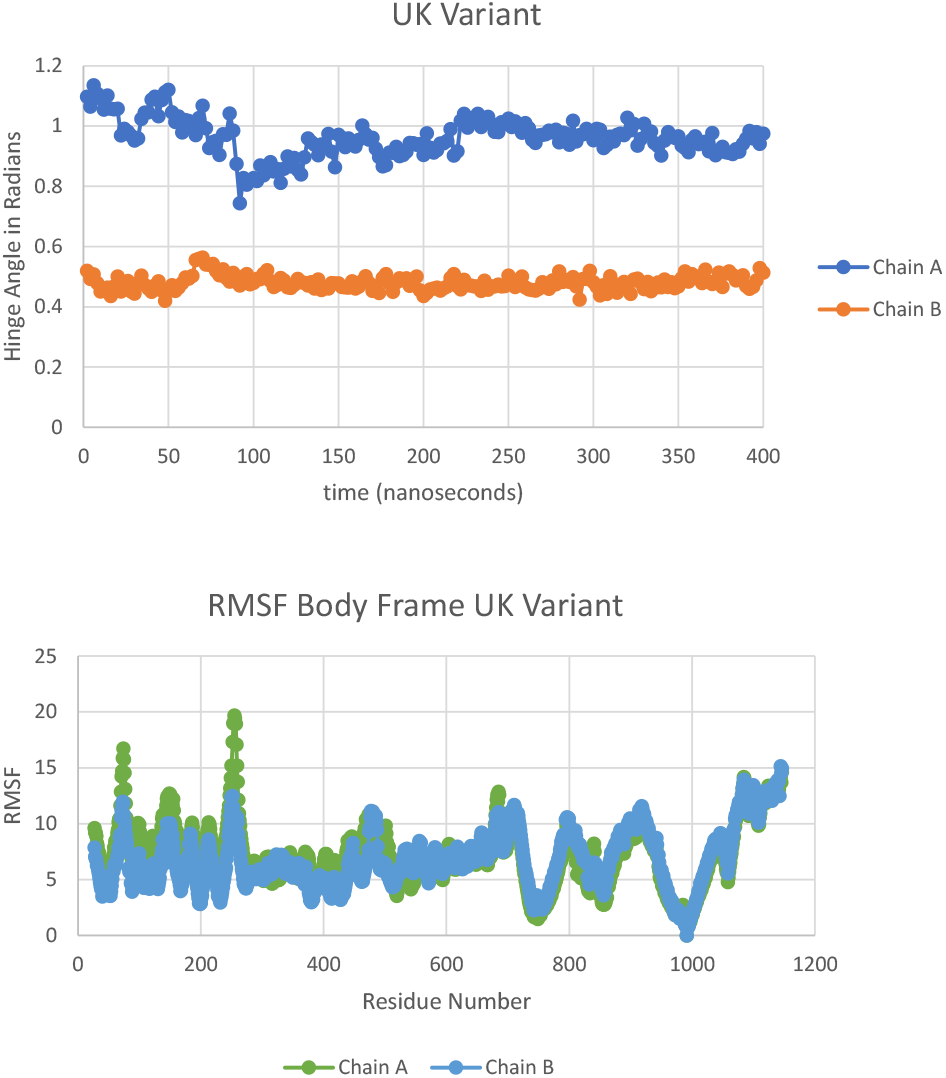
Hinge angle (A) and RMSF values (B) for UK Mutant B.1.1.7.

**Figure 9:**
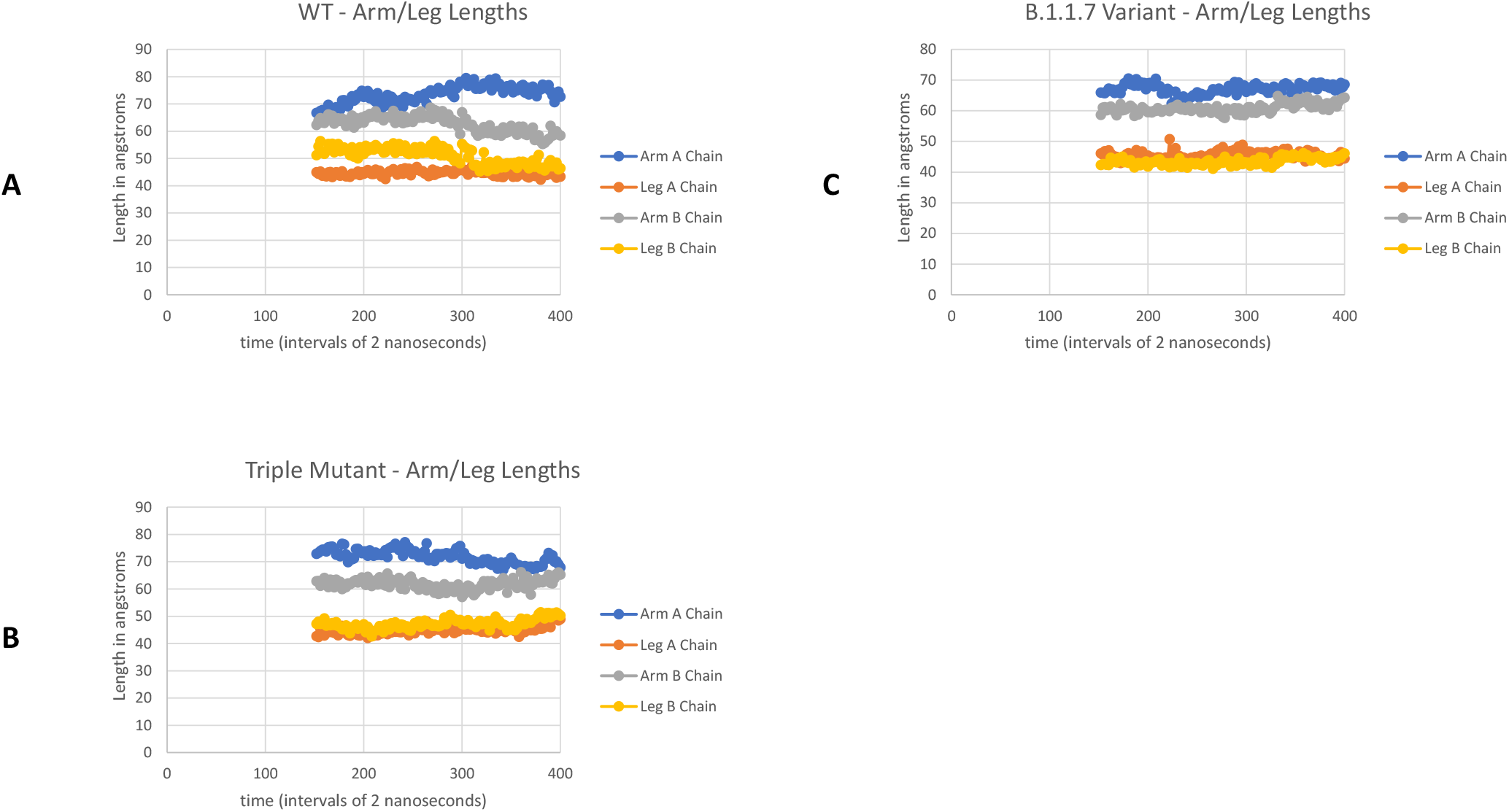
Arm and Leg distance calculations after 150 nsec of simulation for WT, Triple Mutant, and UK Variant B.1.1.7.

## Discussion

Despite its relatively low mutation rate and an inherent error correction mechanism, SARS-CoV-2 continues to display significant numbers of variants due to its high transmission and infection rates. Thus, variants represent a considerable challenge in the current COVID-19 pandemic. The identification of key “glue point” residues that help stabilize the prefusion Spike protein structure could be an important tool in helping to determining structure → function relationships. Here, we have demonstrated the presence of “sequence homologous glue points” by overlaying dominant energetic mappings to sequence alignment maps across lineages of *β* coronaviruses. SARS-CoV-2 prefusion Spike protein was shown to exhibit the least number of stabilizing glue points across these lineages. Additionally, we analyzed a theoretical triple mutant based on the identification of three key stabilizing glue points between neighboring RBD and NTD interactions in the prefusion Spike protein of SARS-CoV-2. We demonstrated the ability to significantly alter protomer configurations by destabilizing these three key residues through alanine mutations or alanine screening. In a reverse sense, by the same methods, we then analyzed the emerging UK Variant B.1.1.7 in order to determine key mutations or deletions in its Spike protein that could potentially be responsible for the more infective and transmissive state of SARS-CoV-2. Our methodologies directly lead to two key mutations D614G and N501Y as possible configuration and ACE2 binding changes, respectively. D614 had been previously identified by us as key glue point associated with dominant energetic interactions between neighboring protomers [11], which in itself demonstrates the potential of glue point monitoring as a helpful tool in tracking possible variants of interest. Biophysical computations demonstrated the configurational changes associated with D614G to a more distal Down state RBD and an overall more stable conformation. We also show that N501Y has a potential hACE2 glue point partner 41Y, which may lead to a strong Y-Y hydrophobic residue pair interaction; this may be partially responsible for the higher infection rate of the UK (B.1.1.7) and SA (B.1.351) variants, although more studies are needed to verify this. It is clear that many tools are needed to quickly translate genome sequence information from WT and variants to potential virion function in order to help direct mitigation strategies and resources in an optimized way, and to ascertain the impact of variants on vaccines and therapeutics.. Here we demonstrate that a combination of protein residue sequence alignment superimposed on glue point and binding point residue identification can flag potential Spike protein function changes and associated viral behavior. These flagged residues can be further analyzed through long-time biophysical computations and experimental methods to more precisely probe functional changes and to assist in the possible identification of variants of interest.

Note Added in Proof: After the initial writing of this manuscript the complete molecular structure of the D614G mutation of the Spike protein of SARS-CoV-2 was determined [25]. Those authors experimentally showed that this mutation did not increase binding through configurational changes from WT, but rather significantly enhanced stability of the prefusion complex. It was argued that the less stable WT configuration leads to premature transitions from prefusion to post fusion complexes. On the other hand, D614G leads to a more stable prefusion state and reduction in premature transitions compared to WT in partial agreement with some of the observations given here. It is not clear, however, if arm length changes observed here have any bearing on the increased transmission and infection rates observed for D614G. Protein configurational changes via mutations/deletions, in general, provide clues to possible changes in transmission which ultimately must be verified experimentally and considered in concert with all aspects of the fusion process.

## Supporting information

ClustalMView.ppt

Table-Headings.txt

Supplemental Movie: UKMutvsWTUpChain.mp4

Supplemental Movie: UKMutvsWTDownChain.mp4

## 5. Acknowledgments

M.H.P. would like to thank the VCU Office of the Vice President for Research, the VCU Department of Chemical and Life Science Engineering, the VCU Center for High Performance Computing, the COVID-19 HPC Consortium (https://covid19-hpc-consortium.org/), and Hoth Therapeutics, Inc. for their continued support of our overall COVID-19 research efforts. O.B. was supported by the National Institutes of Health Institutional and Academic Career Development Award K12GM119955.

## Open Source Software

OpenContact is freely available under the Third-Party Software Tools listings of the Protein Data Bank https://www.rcsb.org or visit http://people.vcu.edu/mpeters

## Competing Interests

V.C.U (M.H.P.) entered an exclusive licensing agreement for “ACE2 Decoy Peptides in the Treatment of COVID-19” with Hoth Therapeutics, NY.

## Supplementary Information

**ClustalMView.ppt**

**Table-Headings.txt**

**Table S1** 6ACD-B-RBDtoA-NTD-finedata.txt

**Table S2** 6CRZ-C-RBDtoA-NTD-finedata.txt

**Table S3** 6M17BchaintoEchain-finedata.txt

**Supplementary Data S1** TripleMutantExtended.xlxs

**Supplementary Movie:UKMutvsWTUpChain.mp4**

**Supplementary Movie:UKMutvsWTDownChain.mp4**

## Notes

### Summary of Updates

This revised manuscript includes computational repeats for improved analysis and trends and a note added in proof on a recently published experimental study of the D614G mutant. Figure 9 on the arm and leg distances associated with the RBD hinge angle has been added. We have corrected that one radian for the hinge angle is on the lower end of the Up state, not the upper end of the Down state, according to the referenced study. Supplementary data on an extended simulation time for the theoretical triple mutant has been added demonstrating the continued transformation from Down to Up.

