## Supplementary figures and images for "Transformations, Lineage Comparisons, and Analysis of Down to Up Protomer States of Variants of the SARS-CoV-2 Prefusion Spike Protein Including the UK Variant B.1.1.7"

### ClustalMView.ppt

#### Slide 1
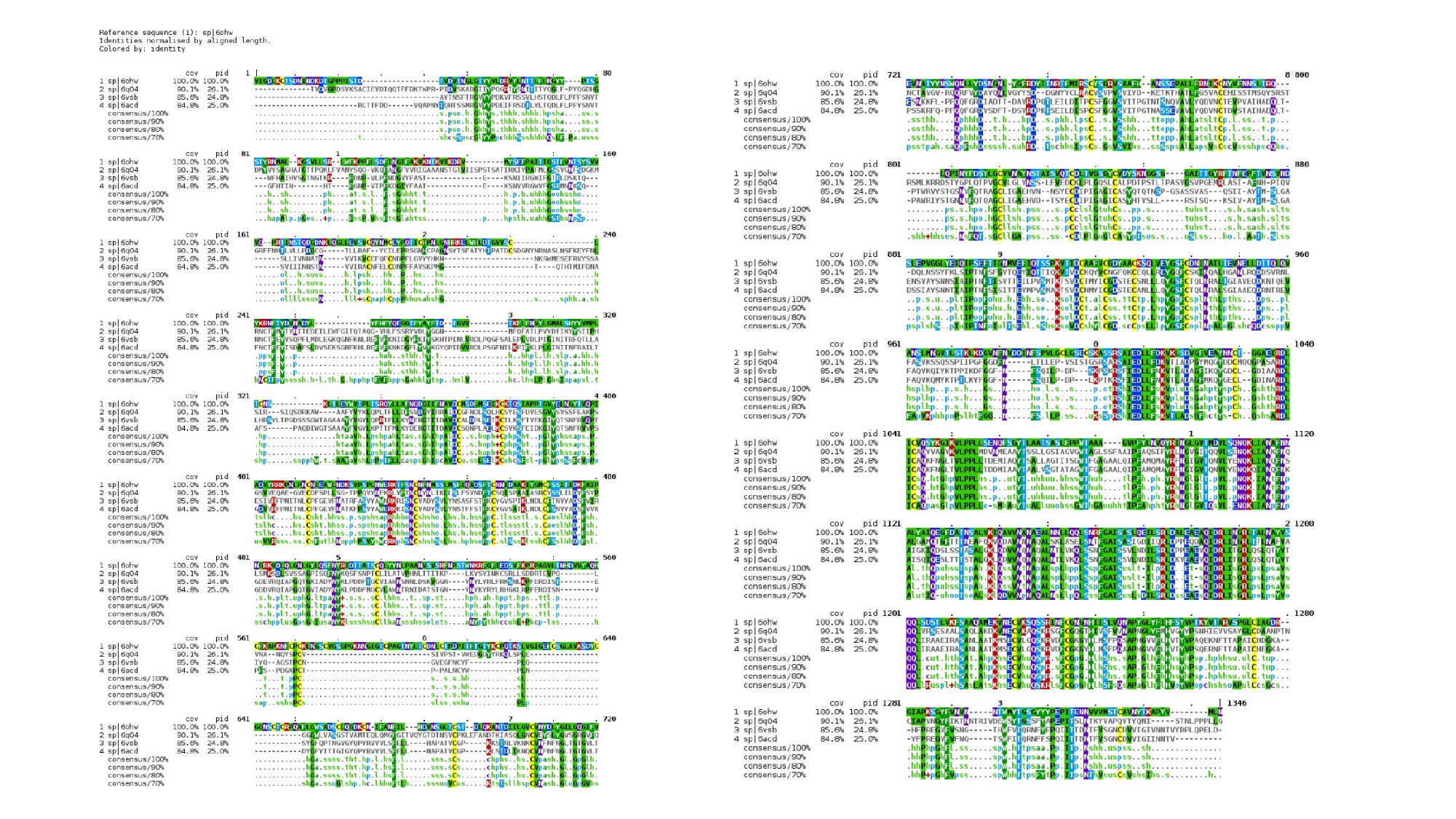
