## Supplementary material for "Transformations, Lineage Comparisons, and Analysis of Down to Up Protomer States of Variants of the SARS-CoV-2 Prefusion Spike Protein Including the UK Variant B.1.1.7": Table-Headings.txt

Table Headings for Supplemmentary Data - From OpenContact: open source code contactgui.f:

Distances rrs are in nanometers
Energies are in kT units
us148 are LJ 6-12 interactions
ucouls are partial atomic charge interactions
All atom force field parameters from AMBER 03


c write fine contact map for matplotlib
c
       write(16,127)resa(j),',',resnma(j),',',atnama(j),',',
     1                      atnuma(j),',', resb(i),',',resnmb(i),',',
     1                      atnamb(i),',',atnumb(i),',',
     1                      rrs,',',us148,',',ucouls
 127   format(a3,a1,i6,a1,a4,a1,i6,a1,a3,a1,i6,a1,a4,a1,i6,a1,e15.8,a1,
     1                      e15.8,a1,e15.8)
	 
	 
when citing OpenContact please use"
 Krall,A., Brunn, J., Kankanala, S., and Peters, M.H. A Simple Contact Mapping Algorithm for Identifying Peptide Mimetics in Protein-Protein Interaction Partners. \emph{Proteins. Structure, Function and Bioinformatics}, 82, 2253-2262, 2014.  DOI:10.1002/prot.24592.
 
 To Download OpenContact go to the PDB website under third party software or visit:
 http://people.vcu.edu/~mpeters
